# Molecular characterization of bacterial communities in sheep cheese through 16S rRNA gene sequencing

**DOI:** 10.1101/753053

**Authors:** Creciana Maria Endres, Ícaro Maia Santos de Castro, Laura Delpino Trevisol, Michele Bertoni Mann, Ana Paula Muterle Varela, Ana Paula Guedes Frazzon, Fabiana Quoos Mayer, Jeverson frazzon

## Abstract

The production of sheep’s milk cheese has grown in recent years since it is a high value-added product with excellent properties. As such, it is necessary to provide data on the microbiota and organoleptic characteristics of this product, as well as the influence of these microorganisms on public health. Thus, the aim of the present study was to characterize the microbial community of different types of sheep cheeses using high-throughput sequencing of the *16S rRNA* gene. The study was conducted with four groups of cheese: colonial, fresh, feta, and pecorino (n = 5 samples per group). The high-throughput *16S rRNA* amplicon sequencing revealed 55 operational taxonomic units in the 20 samples, representing 9 genera of the two bacterial phyla Firmicutes and Proteobacteria. The predominant genera in the samples were *Streptococcus* and *Lactobacillus*. When evaluating alpha diversity by the indexes of Simpson, Chao1, Shannon, and Skew no significant differences were observed between the groups. Evaluating of the beta diversity using Bray-Curtis dissimilarity, the group of colonial cheeses presented a significant difference when compared to the feta (q = 0.030) and pecorino groups (q = 0.030). Additionally, the fresh group differed from the pecorino group (q = 0.030). The unweighted Unifrac distance suggests that the colonial cheese group differed from the others. Moreover, the feta cheese group differed from the fresh group. The distance-weighted Unifrac suggests that no significance exists between the groups. According to this information, the microbiota characterization of these cheese groups was useful in demonstrating the bacterial communities belonging to each group, its effects on processing, elaboration, maturation, and public health.

## Introduction

Sheep cheese production has been increasing globally and represents a major economic driver in many countries, especially in Europe. In Brazil, the main sheep cheeses produced are colonial (locally produced cheese), fresh (not matured), pecorino, and feta. Sheep cheeses are good sources of protein, energy, fat, minerals, and vitamins [1], which have a direct relation to the taste, odor, and texture of the products. Sheep cheeses have a thin, delicate dough, a mild and slightly acidic flavor, thus resembling the typical taste of sheep’s milk. The soft cheeses (fresh and feta) have a relatively high moisture content (44,6% to 45,9%) with a more pronounced acidity, while hard cheeses (colonial and pecorino) have a more solid texture and lower moisture content (35,9% to 38,3%). Apart from these differences in composition, the organoleptic characteristics of these cheeses may vary according to the producing farm, season, animal feeding regimen, and animal breed [2]. Moreover, the microbiota composition also influences cheese characteristics and may have an impact on consumer health. Moreover, the microbial communities of these cheeses represent largely unexplored reservoirs of genetic and metabolic diversity with potential beneficial use in the production of fermented foods.

Classical microbiological techniques have been used to identify lactic acid bacteria as well as deteriorating and pathogenic microorganisms in cheeses. Despite being efficient, these techniques are less accurate and more laborious. Moreover, when a bacterium is not cultivable, it cannot be identified via classical bacteriology. Therefore, methods such as high-throughput sequencing (HTS) based on the *16S rRNA* gene may be more adequate for microbiota evaluation [3]. Thus, the present study aimed to characterize the microbial communities of different types of sheep cheeses and to discuss the presence of microorganisms that are either beneficial or deleterious to human health, including the physiological influence that these microorganisms have on consumers as well as their interference in the sensorial characteristics of the products.

## Materials and methods

### Experimental design and sampling

Four types of sheep cheeses were evaluated in the present study: Feta (n = 5), Pecorino (n = 5), Colonial (n = 5) and Fresh (n = 5). The samples were purchased directly in the company or in supermarkets in Santa Catarina and Rio Grande do Sul states, both in south Brazil. The cheeses were collected on different dates and lots. The samples remained refrigerated until processed in less than 24 h. Sample processing, DNA extraction and MiSeq Illumina sequencing were performed at the Institute of Veterinary Research Desidério Finamor, Diagnostic and Agricultural Research Department, Secretariat of Agriculture, Livestock and Rural Development, Eldorado do Sul, Rio Grande do Sul, Brazil.

Table 1 presents the characteristics of the cheese groups. Fresh and Feta cheese are not ripened, while Colonial and Pecorino are ripened, with lower moisture content. Thus, these cheeses are harder and have a more pronounced flavor.

**Table 1.** Characteristics of cheese groups.

| Group | Moisture Content (%) | Maturation (Days) |
| --- | --- | --- |
| Colonial | 38,3 | 20-30 |
| Feta | 45,9 | 0 |
| Fresh | 44,6 | 0 |
| Pecorino | 35,9 | 45 |

### Sample Processing

Samples of 100 g to 500 g were individually milled in a processor to obtain greater homogeneity. From that, 25 g was withdrawn diluted in 225 mL sterile distilled water and homogenized in a shaker incubator (110 rpm for 2 hours at room temperature). A total of 35 mL of the filtrate was transferred to 50 mL Falcon tube and centrifuged at 10,000 × g for 40 min at 4 °C. A total of 30 mL of the supernatant was discarded and the remainder was homogenized in vortex. One mL of that solution was transferred to a 1.5 mL Eppendorf tube and centrifuged at 14,000 rpm for 5 min at 4 °C for pellet formation.

### DNA extraction

The pellets were suspended in 180 μL of lysis buffer (22.5 mM Tris HCl, 2.5 mM EDTA, 1% Triton X-100, 20 mg/mL lysozyme). The lysis buffer solution was homogenized in vortex and incubated for 1 hour at 37 °C (homogenized every 15 min in the vortex). Subsequently, 20 μL of proteinase K (20 mg/ mL) and 200 μL of Purelink Genomic Lysis/Binding Buffer were added to the samples and incubated at 55 °C for 30 min. Then 200 μL of ethanol (96-100%) were added and the samples were homogenized in vortex for 5 sec. The sample suspension (approximately 640 μL) was used for DNA extraction with Purelink Genomic DNA kit following manufacturers’ instructions. DNA was eluted in 25 uL and stored at −20 ° C until the time of analysis.

### Library preparation and *16S rRNA* sequencing

The 16S PCR libraries were generated amplifying the V4 domain of bacterial *16S rRNA* gene using F515 and R806 primers, both modified to contain an Illumina adapter region [4]. Amplification was performed in a 25 μL mixture, consisting of 12,5 ng of genomic DNA, 1.5mM MgCl_2_, 0.2 μM of each primer, 200 μM of each dNTP, 2 U Platinum Taq DNA Polymerase Platinum (Invitrogen™), and 1X reaction buffer. Amplification was carried out in a BioRad MyCycler Thermocycler (BioRad, USA) according to the following program: initial denaturation at 94°C for 3 min, followed by 30 cycles of 94°C for 30 sec, 55°C for 30 sec, 72°C for 30 sec a final cycle at 72°C for 5 min.

Amplicons were purified using Agencourt AMPure XP beads following manufacturer instructions. Indexes were added to DNA libraries following the manufacturer instructions (Illumina Inc., San Diego, California, USA). Sequencing was conducted on an Illumina MiSeq System with a v2 500- cycles kit.

### Bioinformatics analysis

Bioinformatics analysis of 16S rRNA amplicons were performed with QIIME 2 2019.4 [5]. Raw sequence data were quality filtered and denoised, dereplicated and chimera filtered using the q2-dada2 plugin with DADA2 pipeline [6]. 1,000,000 reads were used for training the DADA2 error model of each sequencing run. The 5’ end 10 nucleotide bases were trimmed from forward and reverse read sequences due to low quality. Reads with a number of expected errors higher than 2 were discarded. Read length filtering was applied and the reads were trimmed at the first instance of a quality score less than or equal to 11. The resulting reads with nucleotide overlap between the forward and reverse reads below 20 and shorter than 240 bp length were discarded. Chimera removal was performed using the “consensus” method, in which chimeras are detected in samples individually, and sequences found chimeric in a sufficient fraction of samples are removed assuming at least 1.0-fold change of potential parents of a sequence being tested as chimeric.

The sequence variants (ASVs) obtained by DADA2 pipeline were merged into a single feature table using the q2-feature-table plugin. All amplicon sequence variants from the merged feature table were clustered into OTU’s with Open Reference Clustering method [7] against the Greengenes version 13_8 with 99% of similarity from OTUs reference sequences [8] using the q2-vsearch plugin with 99% similarity of sequence. The OTU’s were aligned with MAFFT [9] (via q2-alignment) and used to construct a phylogeny with fasttree2 [10] (via q2-phylogeny). Taxonomy was assigned to OTUs using the q2-feature-classifier [11] classify-sklearn naïve Bayes taxonomy classifier. The classifier was trained with extracted Greengenes 13_8 99% OTUs reference sequences truncated at 250 bp length from *16S rRNA* variable region 4 (V4) primer sequences GTGCCAGCMGCCGCGGTAA and GGACTACHVGGGTWTCTAAT for forward and reverse reads, respectively. The resulting feature table from OTU clustering, rooted tree from reconstructed phylogeny, and taxonomy classification were imported from Qiime2 to R v3.6.1 environment using Qiime2R for further data analysis using Microbiome v1.6.0 [12] and Phyloseq v1.28.0 [13] R packages.

### Taxonomic analysis

Feature table was transformed to compositional data for taxa bar_plot composition visualization of the 10 most abundant genera using plot_composition function from Microbiome R package. A heatmap plot was made using NMDS ordination with Bray Curtis dissimilarity using features transformed to log10 frequency using plot_heatmap function from Phyloseq R package.

### Community Diversity Analysis

The OTU’s were aligned with MAFFT [9] (via q2-alignment) and used to construct a phylogeny with fasttree2 [10] (via q2-phylogeny). Alpha diversity metrics (Shannon, Simpson, Chao1 and log_modulo Skew), beta diversity metrics (weighted UniFrac)[14], unweighted UniFrac [15], Jaccard distance, and Bray-Curtis dissimilarity), and Principle Coordinate Analysis (PCoA) were estimated using q2-diversity after samples were rarefied (subsampled without replacement) to 16968 sequences per sample. Alpha diversity significance were estimated with the use of ranks in one-criterion variance analysis [16], well known as Kruskal Wallis test using q2-diversity plugin. Beta diversity significance were estimated using a non-parametric multivariate analysis of variance, well known as PERMANOVA [17].

### Differential abundance Analysis

Feature table was filtered to remove singletons using the q2-feature-table plugin, the OTU’s that was observed less than two samples were removed from the feature table. The resulting filtered features were collapsed at Genus level using q2-taxa plugin for differential abundance analysis. Differential abundance analysis were performed with ANCOM [18] using q2-composition plugin, with mean difference as fold difference in feature abundances across groups and log as transform-function for volcano plot.

### GenBank Accession Numbers

All raw sequence reads have been deposited with the National Biotechnology Information Center (NCBI) and available in BioProject ID SUB6010207.

## Results

### Taxonomic characterization of sheep cheese microbiota

A total of 1,948,389 raw reads were obtained, with 647,832 reads remaining after trimming. A total of 55 OTUs were observed and most (95%) *16S rRNA* gene sequences were classified at the genus level, with only one within the *Enterobacteriaceae* family not being classified. Nine different genera were identified (Fig. 1; Table S4). For a few genera, it was possible to classify microbiota at the species level (Table S1). The number of reads and OTUs are presented in Table S2. Table S3 presents the identified OTUs as well as the frequency at which each OTU was present in the cheese samples.

**Fig 1.**
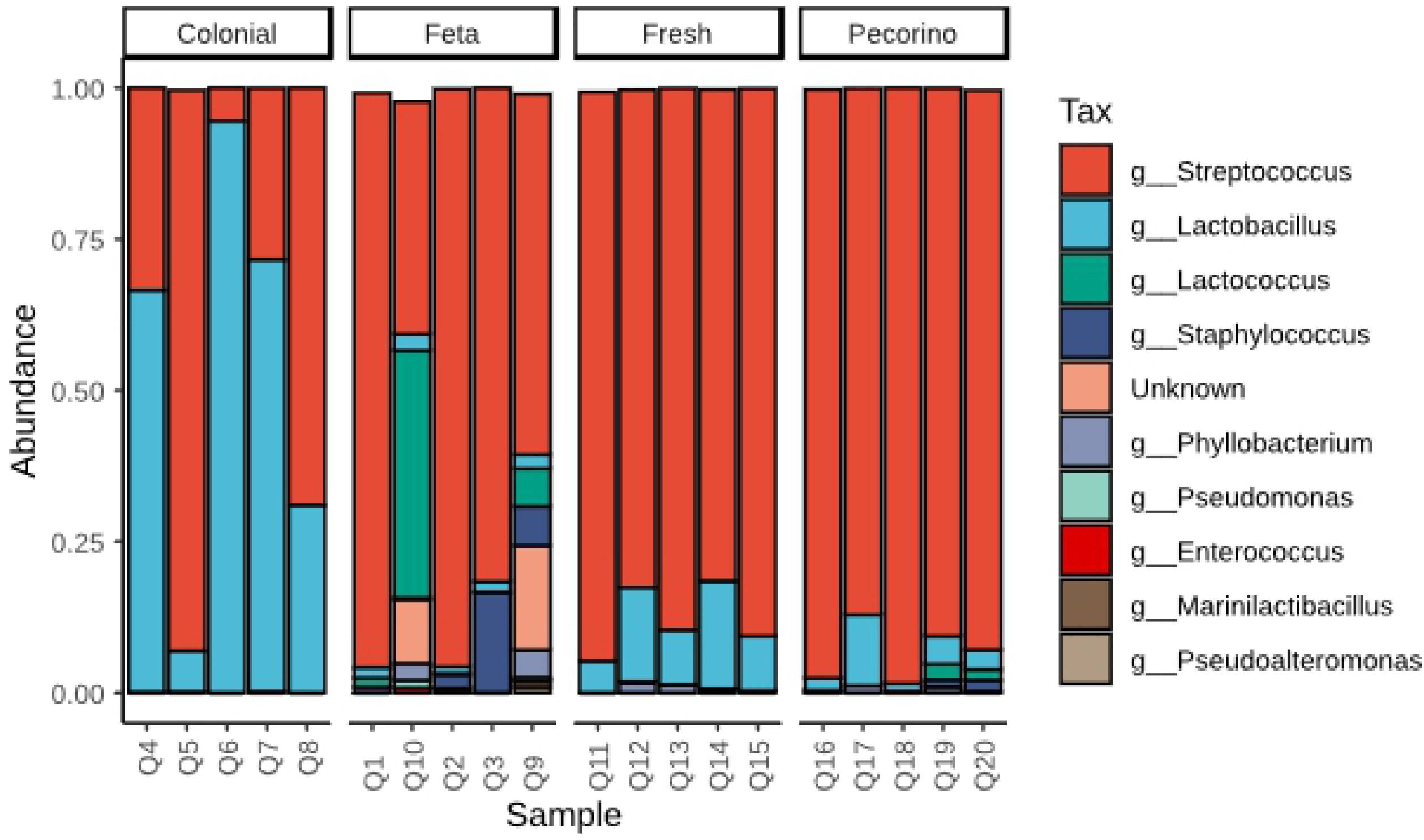
Bar chart detailing the relative abundance of the major genera found on different cheeses.

To determine the percentage by cheese group, see Table S4.

Among the nine observed genera, two core microbiota were present in sheep cheeses. Thermophilic bacteria with lactic acid characteristics prevailed in the cheeses of the fresh group, followed by the pecorino, feta, and colonial groups, with *Streptococcus* being the dominant genus. The genus *Lactobacillus* was also common, as they were identified in all cheese groups (Fig. 2). The presence of genera such as *Staphylococcus*, which may cause disease, was primarily observed in the feta group and at lower frequencies in the fresh and pecorino groups.

**Fig 2.**
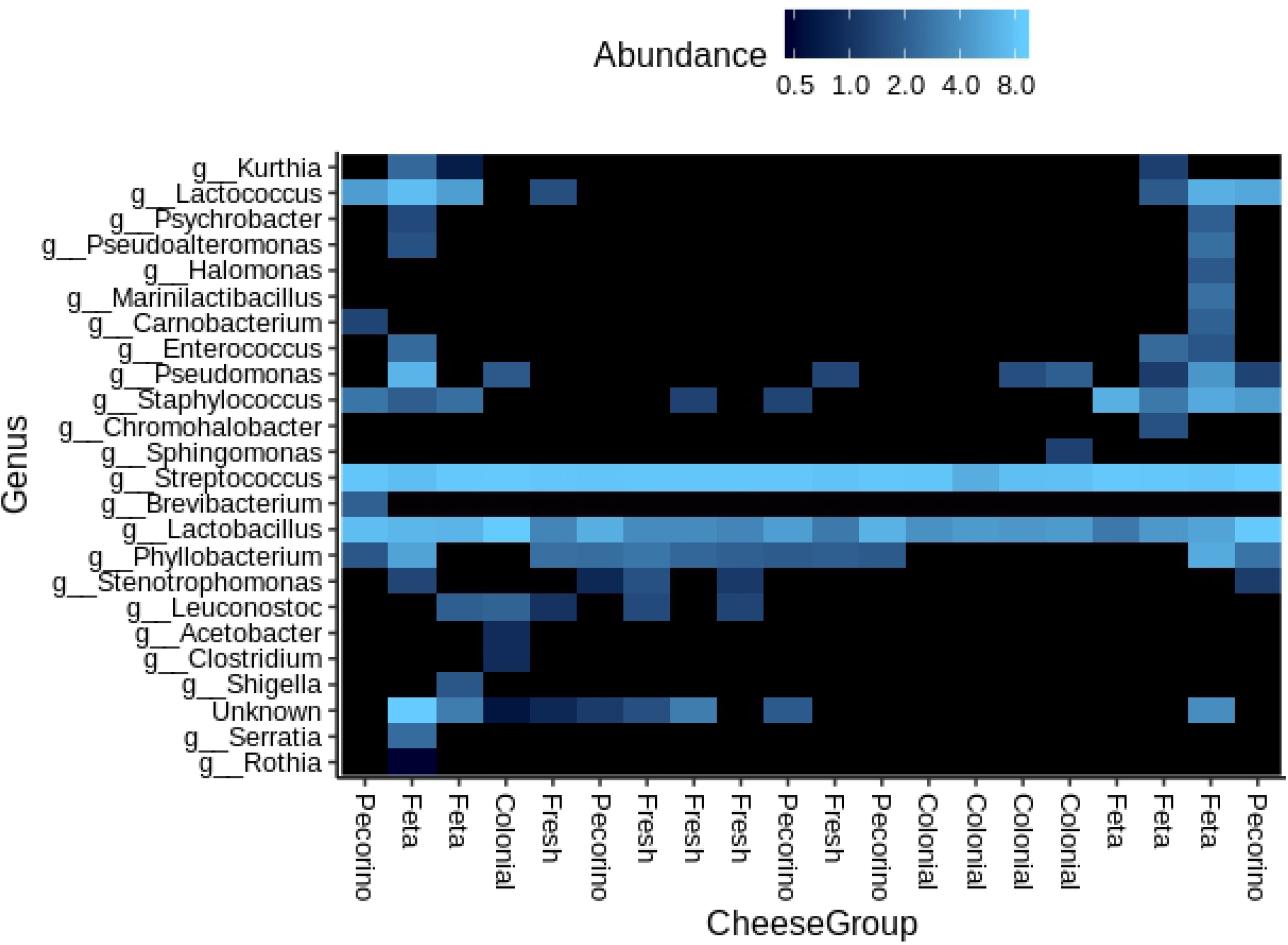
Heat map ordinated using NMDS with Bray-Curtis microbial abundance in cheese samples. Normalization of the frequency of the genera in log_10_. The color scale represents the stepped abundance of each variable, indicated by the score, with blue indicating high abundance and dark blue indicating low abundance.

Upon analyzing the differential composition of the microbiome (ANCOM), of the observed genera, *Lactobacillus* (W = 10) and *Staphylococcus* (w = 8) were differentially observed among the evaluated groups (Fig. 3; Table S5).

**Fig 3.**
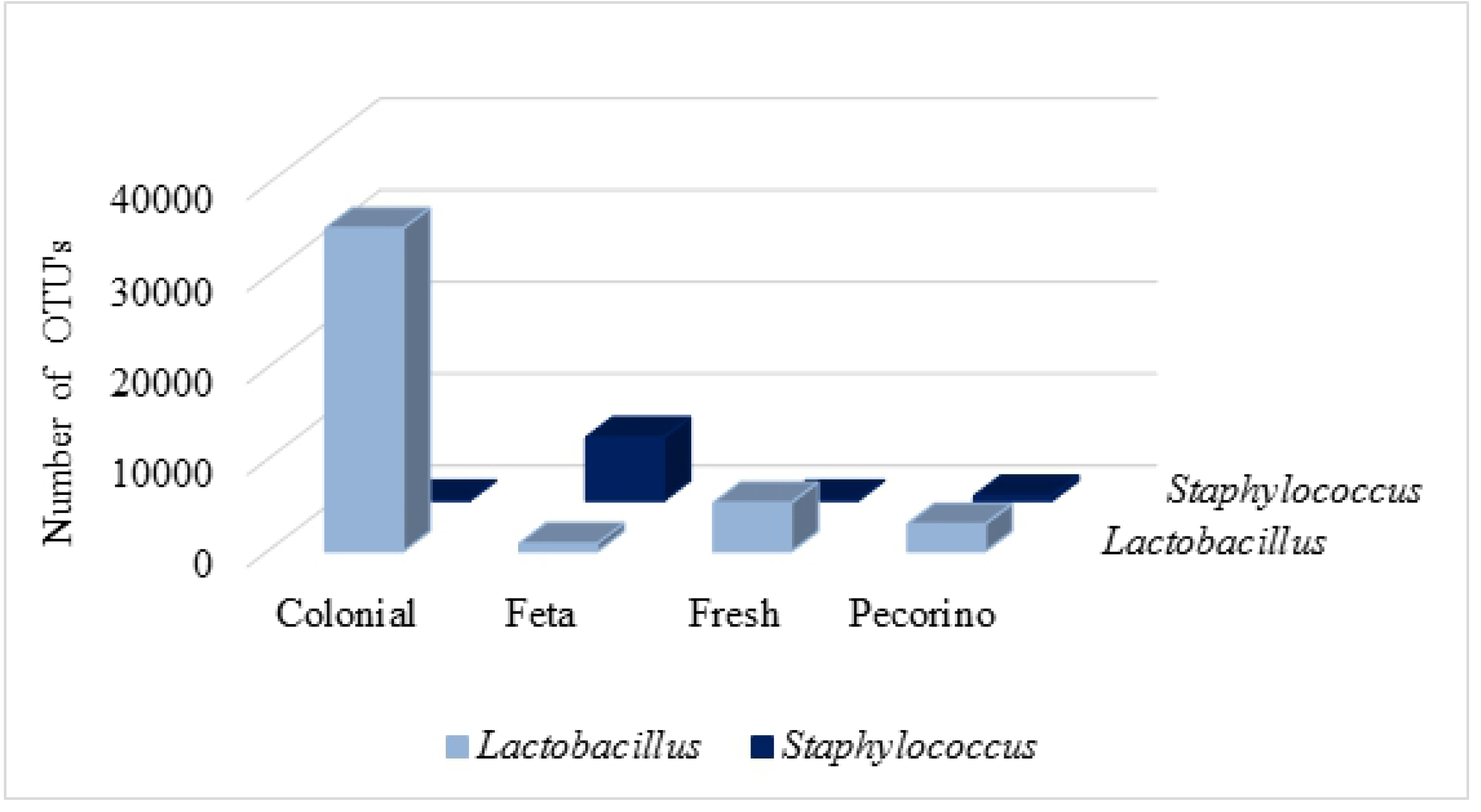
The genera differentially present in evaluated sheep cheese groups.

### Alpha and beta diversity analyzes

There were no significant differences observed in alpha diversity analysis (Fig. 4) compared to Simpson and Shannon indexes. Although the Chao1 index had a p-value of 0.04, no significance was observed among the paired q-values between groups (Table S6).

**Fig 4.**
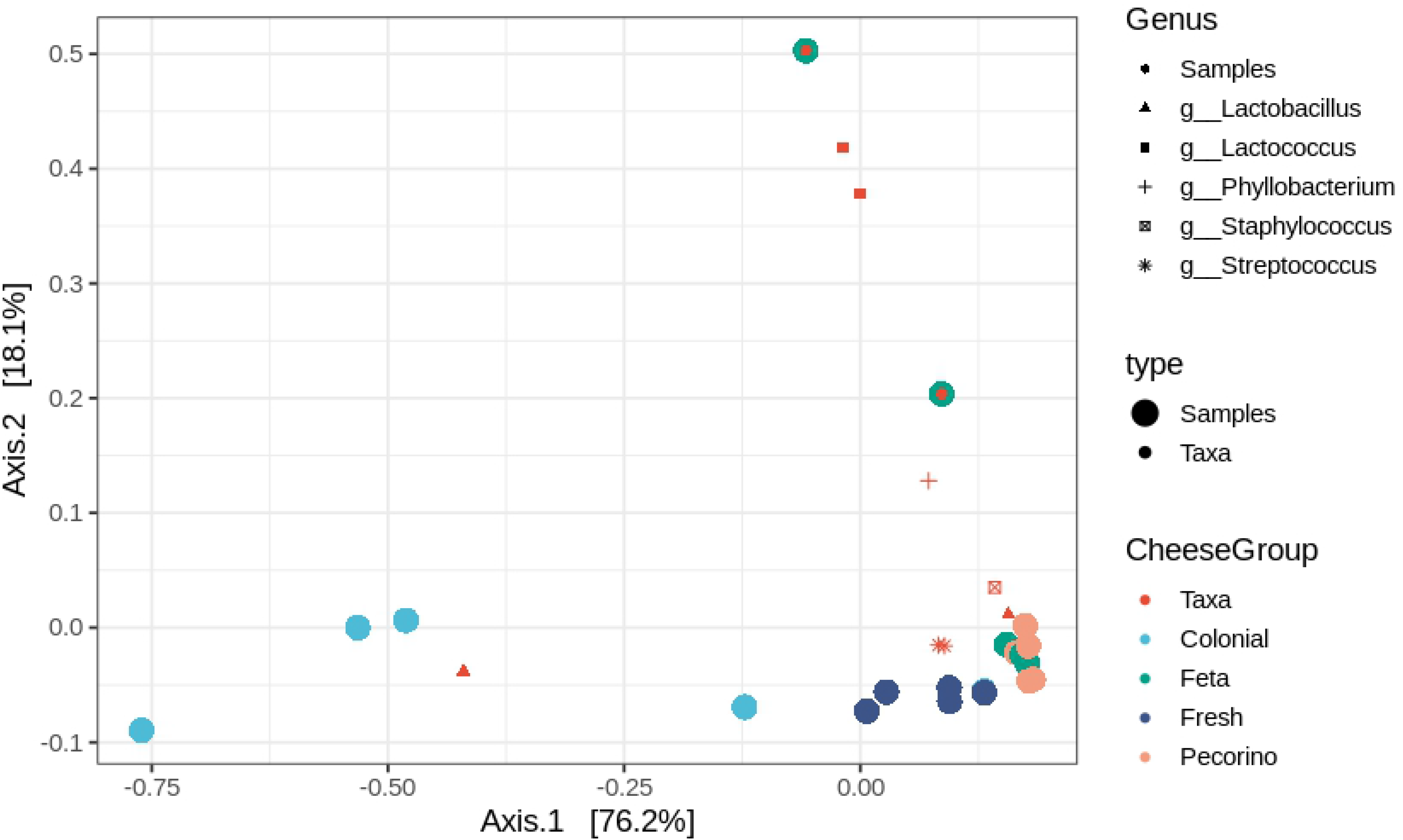
Alpha diversity analysis metrics with Simpson (p-value 0.43), Chao1 (p-value 0.04), Shannon (p-value 0.34), and community rarity index by log_modulo_Skewness (p-value 0.027).

A significant difference was observed in log_modulo_Skewness rarity index between Feta and Colonial cheese groups (0.027) and between Pecorino and Fresh cheese. Based on Shannon and inverse Simpson indexes, the bacterial cheese communities are dominated by a few abundant taxa.

When the Beta diversity (Fig 5) was evaluated by the Bray-Curtis, Unifrac, and Jaccard Index metrics, a significant difference was observed among the groups (Table S7). At Bray-Curtis and Weighted Unifrac, the colonial cheeses were different from feta and pecorino groups showing that bacterial communities in Colonial cheeses have higher dissimilarity and share few taxa compared to other Cheese Groups. The Unweighted-Unifrac distance showed that all the cheese groups are phylogenetically distant between them, having only few taxes closely related. At Jaccard distance, all cheese groups are different, except for feta and pecorino group.

**Fig 5.**
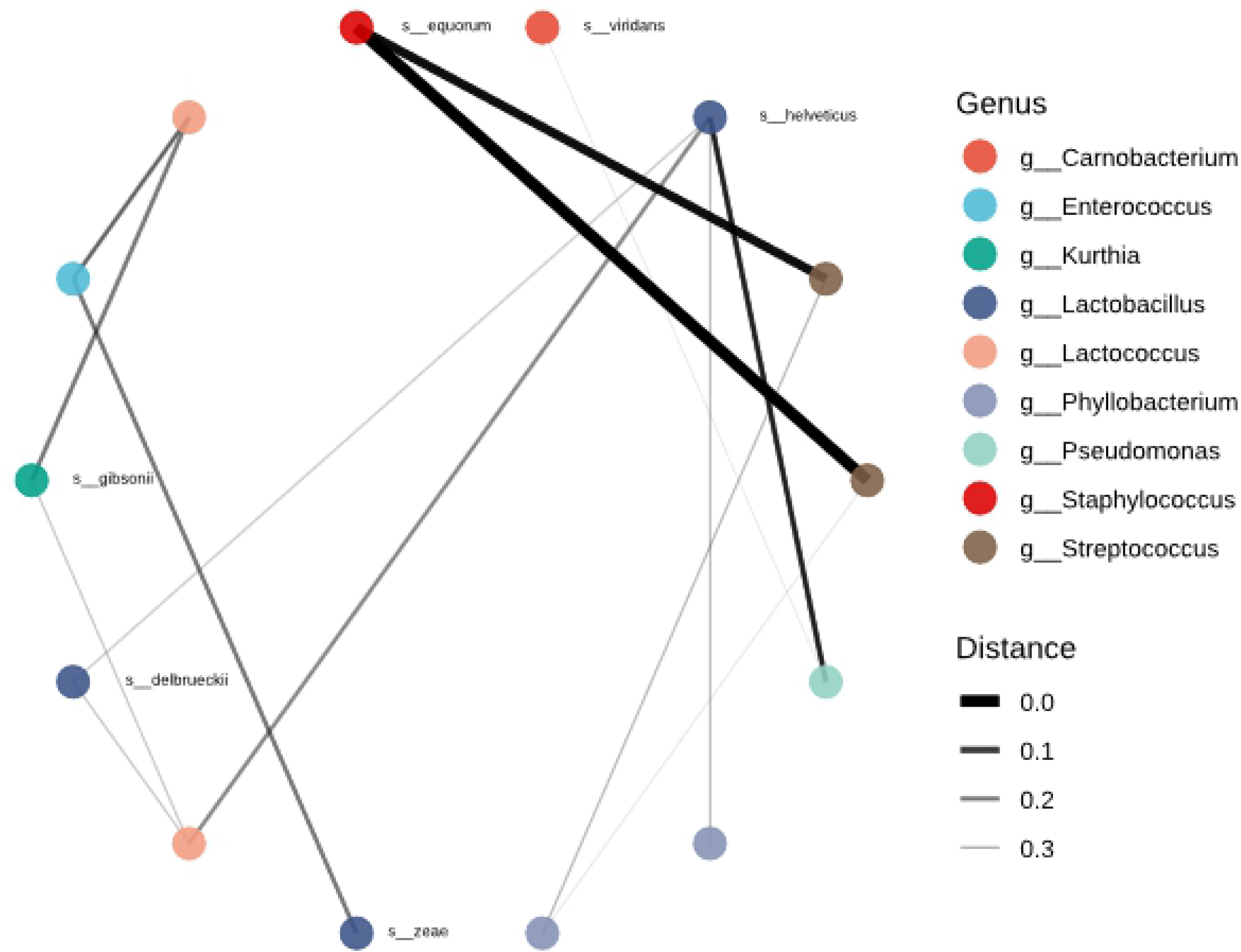
Beta diversity PCoA plots for cheese groups. A significant difference was observed among groups evaluated using Bray-Curtis dissimilarity (p-value 0.0014), weighted Unifrac (p-value 0.0006), unweighted-Unifrac (p-value 0.0001), and Jaccard index (p-value 0.0001).

A PCoA biplot analysis (Fig 6) was created to observe the relationship between features collapsed at the genera level with the variance of plots. A similar pattern was observed for both Bray-Curtis dissimilarity and weighted Unifrac, demonstrating that *Lactobacillus* and *Lactococcus* are the main genera contributing to diversity.

**Fig 6.**
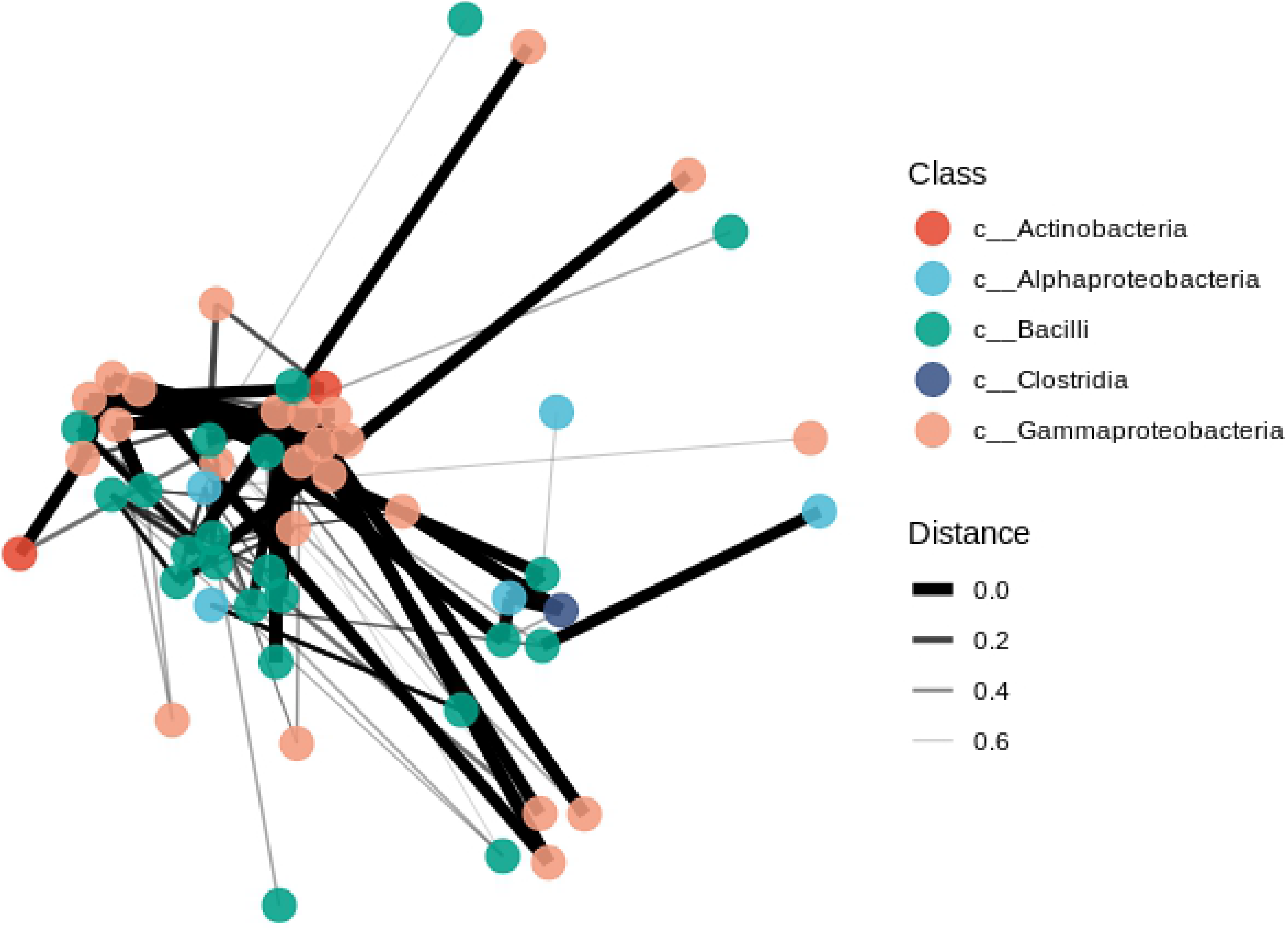
PCoA Bray-Curtis dissimilarity biplot by cheese group at the genus level.

A network was inferred using Bray-Curtis distances to compute node connections at genus level collapsed at species level when detected. The network showed that *Lactobacillus helveticos* are one of the main species, closely abundant related to other genus in all cheese samples. *Staphyloccocus equorum* were the species most closely related to *Streptococcus* (Fig 7). The Fig 8 presents the relevance between the classes of bacteria present in the cheese.

**Fig 7.**
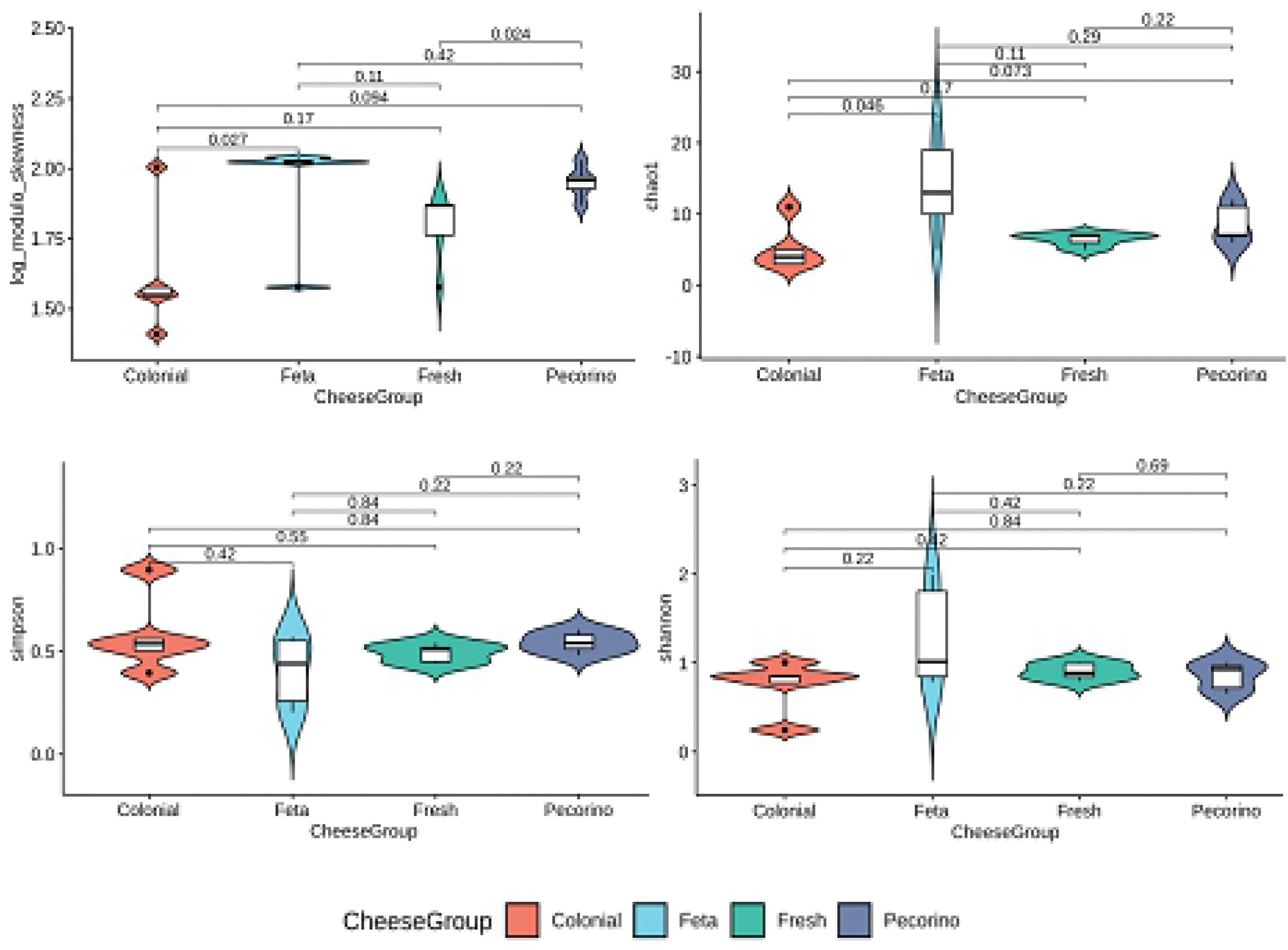
Network inference of the core microbiome using Bray-Curtis distance at the genus level showing species interactions.

**Fig 8.**
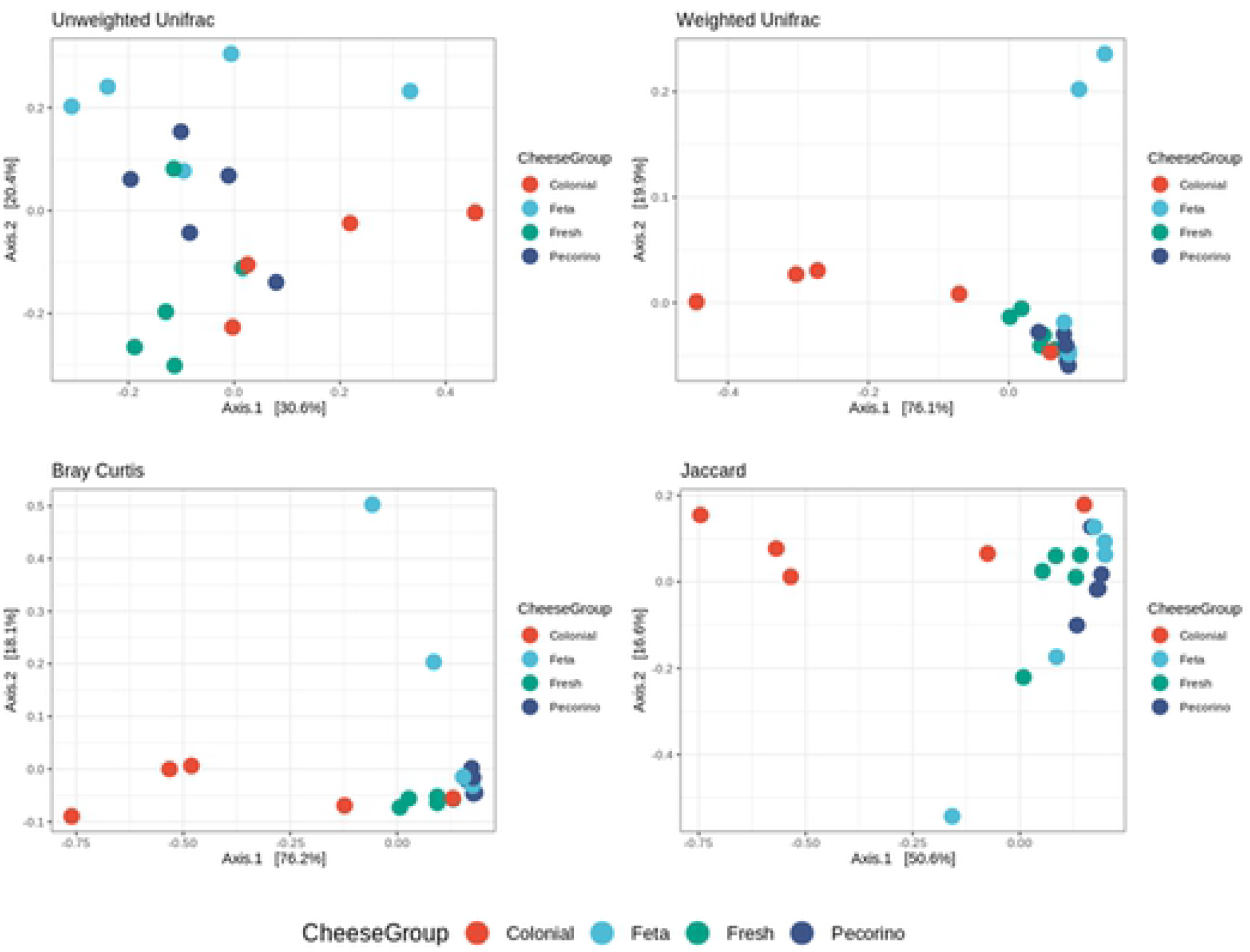
Network inference of microbiome using Bray-Curtis distance at the class level.

## Discussion

Microbiota are a relevant factor in the area of food, being necessary to characterize it in different products since it is responsible for all sensory characteristics and product quality. Furthermore, microbiota may be beneficial in the case of probiotic microorganisms or harmful in the case of spoilage and pathogenic microorganisms. While the microbiota has been studied for various foods, further clarification is required. One prominent product is sheep’s cheese since it includes raw material milk that is inherently composed of nutrients for the development of various microorganisms. In addition, sheep cheese is a product that is increasing in production, making it necessary to thoroughly understand the microbiota within it.

The characterization of sheep cheese microbiota was performed in this study, presenting the differences in microbiota composition associated with different types of cheeses and moisture contents. Through the characterization of cheese microbiota, it is possible to evaluate the presence of predominant genera in cheese and to associate their effects on product characteristics as well as the impact that certain microorganisms present for public health. As expected, the genera *Streptococcus, Lactobacillus*, and *Lactococcus* were identified as predominant, since the starter cultures used in the cheese production process are generally composed of *Streptococcus thermophilus, Lactobacillus bulgaricus, Lactobacillus helveticos*, and *Lactobacillus casei* strains. These microorganisms contribute to the development of the taste and texture of cheese, while some species produce bacteriocins that contribute to their conservation by inhibiting the growth of pathogenic and spoilage microorganisms. A predominant microbial composition of lactic acid bacteria is expected in cheese [19].

Studies performed [20] using *16S rRNA* gene sequencing revealed very simple microbial communities in mozzarella, grana padano, and parmigiano reggiano identified *Lactobacillus* as the predominant genus. The genera *Streptococcus* and *Lactobacillus* were identified as predominant in different types of cheese made from cow and buffalo milk, which is corroborated by the present study [21–23]. *Lactobacillus* species have the capacity to develop in low pH and high salt concentrations (i.e. the environment of cheeses). Due to this capacity, it has already been reported as abundant in cheeses. *Streptococcus* was described as the dominant genus in the Cerrado, while the genera *Lactobacillus*, *Lactococcus*, and *Leuconostoc* were also reported in Canastra cheese [24]. *Streptococcus* have a great influence on the texture, taste, and aroma of cheese products, while also producing bacteriocins that protect the product from microbial spoilage [25].

*Lactobacillus* sp. grow at a very low rate during the initial weeks of cheese ripening but may dominate the cheese microbiota following the initial culture death phase. These bacteria are important both in the processing of fresh and ripened cheeses during the ripening process, as they release bioactive peptides, vitamins, and oligosaccharides [26]. This explains the predominance of *Lactobacillus* sp. in colonial cheese as well as the low abundance in fresh products. Notably, in cheese with longer maturation periods, these microorganisms may die due to low pH and high salt concentrations [27]. The *Lactococcus* identified in the present study have already been described in cheese and yogurt, thus *Lactobacillus* sp.*, Streptococcus* sp. and *Lactococcus* sp. can be defined as the central cheese microbiota. Other observed genera in evaluated cheeses include *Enterococcus* sp.*, Streptococcus* sp., and *Leuconostoc* sp., which were also described in cow, sheep, and goat milk and cheese. Additionally, *Phylobacterium* sp.*, Carnobacterium* sp.*, Stenotrophomonas* sp.*, Pseudomonas* sp., and *Kurthia* sp. were also detected [28].

In addition to the beneficial microbiota, there may be genera involved in spoilage and pathogenic microorganisms, such as *Staphylococcus* sp. and *Enterobacteriaceae* sp. Moreover, the high abundance of *Staphylococcus* sp. is related to cheese processing problems such as pasteurization process failures, improper equipment hygiene, manipulators, and incorrect storage temperatures. The abundance of *Staphylococcus* sp. observed in the feta cheese group in this study can perhaps be associated with these problems. Moreover, the presence of *Staphylococcus* sp. in feta cheese has also been reported by other authors. The genus *Pseudomonas* has been described in spinach, raw meat, and raw milk, while they are also recognized as common agents that contribute to the deterioration of several types of food [29, 30].

Alpha diversity evaluated by the Simpson, Shannon, and Chao1 metrics did not present significant differences. Upon evaluating the heat map, lower diversity was observed in the colonial cheese group, which may be associated with the quality of the raw milk and the high abundance of L*actobacillus* sp. and *Streptococcus* sp. microorganisms that can inhibit the growth of others. However, upon evaluating beta diversity by Bray-Curtis dissimilarity, unweighted Unifrac, weighted Unifrac, and Jaccard index measurements, a significant difference was observed between some groups. This difference was verified when analyzing the q-value, since assessing only the p-value is not sufficient. However, q-value is also considered. This is because q-value is a p-value that has been set to the false detection rate (FDR). The FDR is the proportion of false positives that can be expected when conducting a test. In contrast to the q-value, the p-value provides the probability of a false positive in a single test. In the case of metagenomics, the q-value must be used since we performed several tests for a sample.

Significant differences in beta diversity are usually associated with the use of different raw milk, i.e. from different regions and producers [29]. In the study region, the cheeses (colonial, fresh, feta, and pecorino) are made from pasteurized sheep’s milk and use starter cultures in their production. The primary difference is the maturation period and moisture content of the samples. Authors who studied the microbiota of cheeses made from raw milk and pasteurized milk observed high biodiversity and statistical difference between these groups; according to the author, this difference is directly associated with the pasteurization process, with cheese made from raw milk showing more diversity [31].

In the present study, different methods were used to access the beta diversity, with divergent results among them. The Jaccard and Unweighted Unifrac coefficients measure the distance between communities based on the species they contain and does not take into account abundance, being qualitative metrics. In the context of our cheese groups, Jaccard and Unweighted Unifrac showed higher biodiversity difference. These results were related to the observed by Delcenserie et al [31]. On the other side, the Bray-Curtis and Weighted Unifrac distances take abundance into account. Bray-Curtis showed that pecorino, fresh and feta cheeses samples have similar communities and share higher abundant taxa. Weighted Unifrac showed that abundant taxa like *Streptococcus* and *Lactobacillus* are phylogenetically closed in different cheese communities. Thus, related that previous studies showing that these two genera are the main starter cultures and wide abundant in cheese samples.

The observed results show variation in microbiota depending on cheese type, thereby suggesting that variations in processing, storage, and ripening conditions are important factors in shaping the cheese microbiota. Although lactic acid bacteria were predominant, it is necessary to elucidate the microbiome diversity of sheep cheese samples since the production of this type of product is increasing. Moreover, it is important to note that this diversity could be related to processing, hygienic habits during processing, and the raw materials and ingredients used, which are directly related to the characteristics of the cheese. The microbiome result can readily be used to improve processing conditions and product quality while additionally facilitating the possible standardization of legislation for dairy products of sheep origin.

## Conclusions

The present study demonstrated that the characterization of the microbiota communities of different groups of sheep cheeses was useful for subsidizing data on the bacteria present, thereby evaluating their influence on the composition and microbiological quality of sheep cheeses, something that has been largely absent from the literature to date. The use of metagenomic analysis allowed for the identification of different bacterial genera in the four studied cheese groups.

## Acknowledgements

JF is grateful for CNPq for Financial support (#401714/2016-0 and # 305495/2018-0).

